# Metabolic engineering approach to boost γ-decalactone production in *S. cerevisiae* identifies a critical role for the peroxisomal Fatty Acyl-CoA Synthetase Faa2

**DOI:** 10.1101/2025.07.21.665937

**Authors:** David Boy Chia, Jeng Yeong Chow, Mohammad Alfatah, Shawn Hoon, Prakash Arumugam, Hong Hwa Lim, Uttam Surana

**Affiliations:** Institute of Molecular and Cell Biology (IMCB), Agency for Science, Technology and Research (A*STAR), 61 Biopolis Drive, Proteos, Singapore 138673 Singapore; Bioinformatics Institute (BII), Agency for Science, Technology and Research (A*STAR), 30 Biopolis Street #07-01 Matrix Singapore 138671, Singapore; Singapore Institute of Food and Biotechnology Innovation (SIFBI), Agency for Science, Technology and Research (A*STAR), 31 Biopolis Way, Nanos, Singapore 138669, Singapore; School of Biological Sciences, Nanyang Technological University, Singapore; Singapore Immunology Network (SIgN), Agency for Science, Technology and Research (A*STAR), 8A Biomedical Grove, #04-06, Immunos, Singapore 138648, Singapore; Department of Pharmacology, National University of Singapore, Singapore

## Abstract

Lactones constitute a family of aroma compounds found in fruits, flowers and vegetables and are in high demand in the food industry. Previous studies have reported biotransformation of castor beans-extracted ricinoleic acid to γ-decalactone using oleaginous yeast *Yarrowia lipolytica*. Given the potential toxicities associated with castor beans, we have used a metabolic flux-engineering approach to produce γ-decalactone from oleic acid in *Saccharomyces cerevisiae*. Intracellular conversion of oleic acid to ricinoleic acid was achieved by the expression of oleate hydroxylase Fah12 from ergot fungus *Claviceps purpurea*. Glycerol-3-phosphate dehydrogenase-mediated glycerol synthesis was identified as the major metabolic diversion of oleic acid that negatively impacts γ-decalactone yields. Chemogenomic profiling analysis revealed that the tryptophan biosynthetic pathway provides resistance to γ-decalactone-mediated toxicity in yeast. Overexpression of tryptophan transporter Tat1 enhanced γ-decalactone production by about 3- fold. Deficiency of genes encoding the cytoplasmic fatty acyl CoA synthetases *FAA1* or *FAA4* alone did not significantly influence γ-decalactone production. However, deficiency of peroxisomal *FAA2* drastically diminished the yield of γ-decalactone. Thus, this study uncovers the metabolic barriers to oleic acid-to-γ-decalactone conversion and identifies Faa2 as an essential element in this biotransformation.

## Introduction

Bioflavour and fragrance compounds are extensively used by cosmetic and food/nutrition industries to augment the appeal of their products. The market value of flavor and fragrance compounds has continued to rise steadily over the last couple of decades and is expected to reach USD 16.1 billion by 2027 (1). Extraction of aroma compounds from fruits and plants is an expensive and a difficult task. For instance, the high demand for natural 2-phenylethanol or vanillin was not met due to expensive extraction/purification processes and low yields (2, 3). Moreover, agricultural factors such as weather, seasons and sensitivity of crops to plant pathogens also affect production costs. Currently, aroma compounds are mainly produced through chemical synthesis. However, these processes involve use of organic solvents and environment-unfriendly methods that yield racemic mixtures of the desired compound. Therefore, biotechnological methods involving use of microbial hosts represent an attractive and efficient approach for sustainable production of non-synthetic bioflavours.

Aroma compounds belong to different chemical classes such as ketones, esters, alcohols, aldehydes, or lactones (4). Of these, lactones are a ubiquitous group of cyclic esters with notable odors of peach, coconut or strawberry. Because of their distinctive aroma, γ- and ẟ-lactones have been the focus of many microbial production processes. γ-decalactone is perhaps the most valuable because of its fruity and floral aroma with low perception threshold. Biotransformation of hydroxy fatty acids through β-oxidation pathway is a key process for the microbiological production of lactones (5). Some microorganisms such as *Trichoderma viride*, *Sporidiobolus salmonicolor*, *Fusarium poae* and *Ashbya gossypii* can carry out de novo biosynthesis of lactones (6). The oleaginous yeast *Yarrowia lipolytica* has also been described to efficiently convert ricinoleic acid (12-hydroxy-cis-9-octadecenoic acid) from castor oil to γ-decalactone with a conversion yield of 22.7% (w/w) (7). The oleaginous yeast *Waltomyces lipofer* was found to be even more efficient in the degradation of 60g/L 12-Hydroxystearic acid (produced by hydrogenation of castor oil) to 27.6 g/L γ-decalactone (conversion yield = 26%) (8). Both approaches are dependent on the availability of castor beans which are considered unsuitable due to potential toxicities associated with them (9). Use of oleaginous yeasts is also challenging due to their propensity to degrade ricinoleic acid to several different substrates that leads to accumulation of other lactones like 3-hydroxy-γ- decalactone, 2-decen-4-olide or 3-decen-4-olide, thus resulting in impurity of the desired product (10). An alternative to castor seeds is the production of hydroxy fatty acid, such as ricinoleic acid, by the microorganism used for lactone production. Such studies were first performed using *Pseudomonas* strains for biotransformation of oleic acid to 10-Hydroxystearic acid (11). Subsequently, the production of ricinoleic acid in *Pichia pastoris* was achieved with the hydroxylase CpFAH and acyl-CoA dependent diacylglycerol acyltransferase CpDGAT from the fungus *Claviceps purpurea* (12). The disadvantage of using *P. pastoris* in this context is the presence of Δ12-desaturases, which compete with the hydroxylase for oleic acid (12).

*Saccharomyces cerevisiae* is a GRAS (Generally Regarded as Safe) microorganism with a comprehensively characterised fatty acid synthesis/degradation pathways and a single desaturase. It has been used in a wide range of industrial-scale fermentation processes to produce various chemicals (13). Hence, it can serve as an appropriate host to produce γ-decalactone independently of castor oil to maintain the precursor/product purity. Oleic acid is nontoxic and an easily available resource to serve as a precursor for lactone synthesis. However, *S. cerevisiae* lacks endogenous hydratase or hydroxylase to convert long chain fatty acids (saturated and unsaturated) to the hydroxyl fatty acid which can be subsequently metabolised through cycles of β-oxidation to yield γ-decalactone. In this study, we use an imported hydroxylase in combination with metabolic flux engineering strategies to demonstrate the metabolic capacity of *S. cerevisiae* for γ-decalactone synthesis. While seeking to define metabolic barriers to γ-decalactone production, we have uncovered glycerol synthesis pathway as the most significant metabolic diversion of oleic acid. Chemogenomic profiling analyses showed that genes involved in tryptophan biosynthesis and transport provide resistance to γ-decalactone-mediated toxicity. Unexpectedly, peroxisomal fatty acyl CoA synthetase Faa2 emerges as a critical enzyme for biotransformation of oleic acid to γ- decalactone. Our findings have important implications for the metabolism of hydroxy fatty acids by *S. cerevisiae* and establishes a platform for synthetic biology-guided production of flavour lactones.

## Results

### Identification of *C. purpurea* hydroxylase Fah12 for hydroxyl fatty acid synthesis in *S. cerevisiae*

To demonstrate the natural capacity of *S. cerevisiae* to metabolise ricinoleic acid (12-hydroxy-cis- 9-octadecenoic acid) to γ-decalactone, WT BY4741 strain was cultivated in the YEPD medium containing 1 mM (289 mg/ml) ricinoleic acid for 24, 48 and 72 h. The presence of γ-decalactone in the medium was analyzed at different time points using GC/MS. The WT BY4741 exhibited maximum yield of ∼70 mg/L after 72 h of cultivation (Figure 1A). While *S. cerevisiae* can convert ricinoleic acid to γ-decalactone, it lacks the enzyme to convert oleic acid to hydroxy fatty acid. To search for enzymes that would capacitate *S. cerevisiae* to convert oleic acid to hydroxy fatty acid, six genes encoding C10-hydratase, one encoding C13-hydratase and one encoding C12- hydroxylase were imported from other organisms (Table 1), codon-optimised and tagged with HA3 epitope. All eight genes were cloned under the control of *ADH2* promoter and integrated at the *Δ15* locus of the *S. cerevisiae* genome. Expression of these genes was verified by Western blot analysis (data not shown). Cells were cultivated in YEPD medium for 72 h and the presence of hydroxy fatty acids was analyzed using GC/MS. While oleate hydratases TiOLH, AlOLH and NuOLH yielded no hydroxy fatty acids (data not shown), FAH12 from *C. purpurea* produced 12- hydroxy-cis-9-octadecenoic acid (ricinoleic acid) from oleic acid (Figure 1B &1C). The EmOHYA from *Elizabethkingia meningoseptica* was able to convert oleic acid and linoleic acid to 10-Hydroxystearic acid and 10-Hydroxy-12-octadecanoic acid, respectively. The enzymes SmOLH and LfOLH showed oleate hydratase activity (Figure 1B & 1C). As previously reported, LaOLH from *Lysinibacillus acidophilus* exhibited conversion of linoleic acid to 13-hydroxy-9- octadecenoic acid (Figure 1B & 1C). The enzymes with proven activity towards oleic acid were then tested *in vivo* for the production of respective lactones derived by degradation of the hydroxylated fatty acids. γ-decalactone and γ-dodecalactone could be detected in the culture supernatant only for CpFAH12 and EmOHYA, respectively (Figure 1B-C).

**Figure 1.**
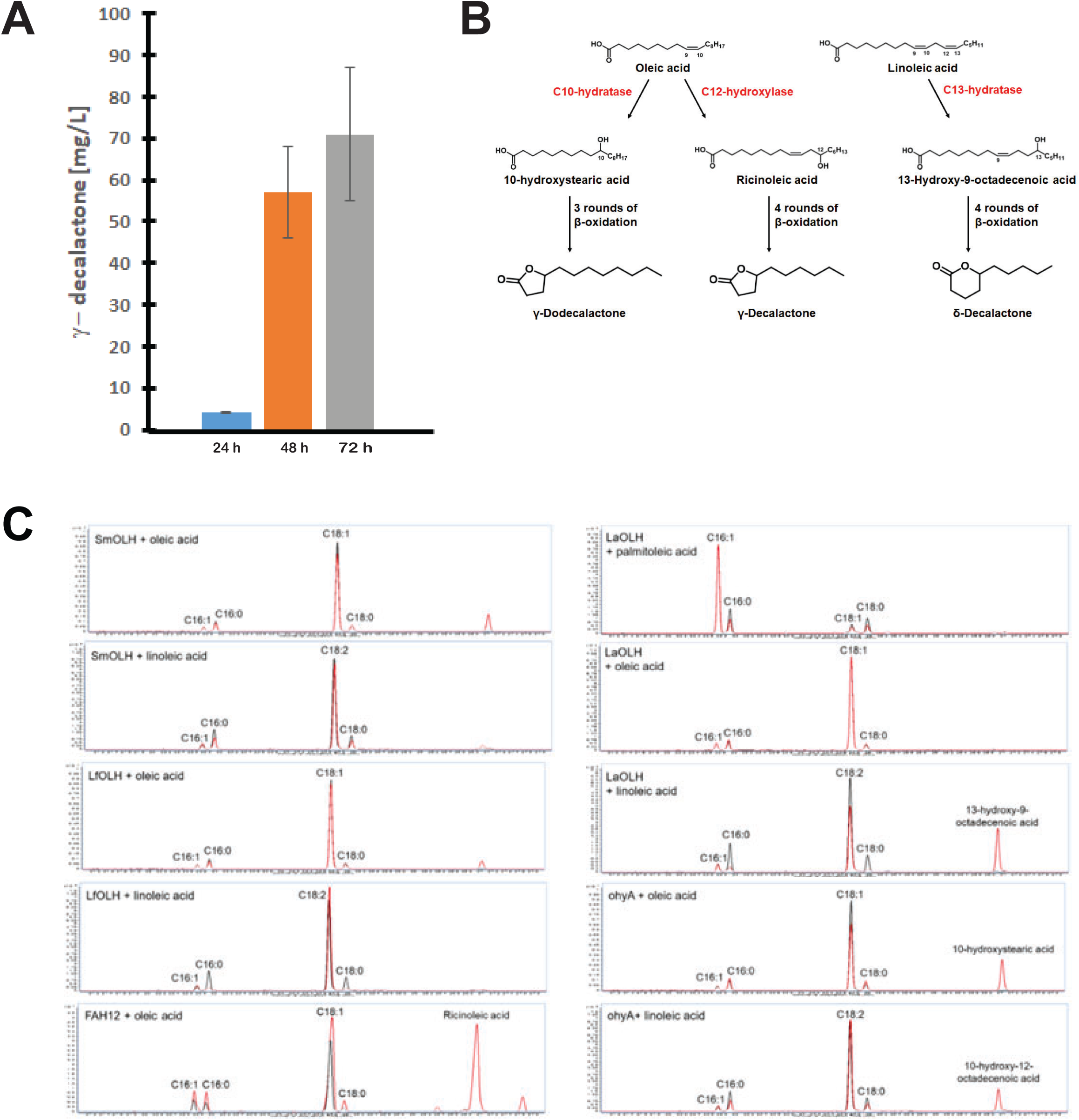
**Production of lactones from fatty acids in yeast** A) WT BY4741 strain was cultivated in the YEPD medium containing 1mM (289 mg/ml) ricinoleic acid for 24, 48 and 72h. The presence of γ-decalactone in the medium was analyzed at different time points using GC/MS B) Expected reaction products with expression of C10-hydratase/C-12 hydroxylase and C13- hydratase with yeast cultures with oleic acid and linoleic acid substrates respectively C) Fatty acid hydratases and hydroxylases were expressed under the control of the *GAL1-10* promoter. Cell lysate was incubated with 1mM of one specific unsaturated fatty acid (palmitoleic acid, oleic acid, linoleic acid) and the products were detected by GC-MS.

**Table 1.**
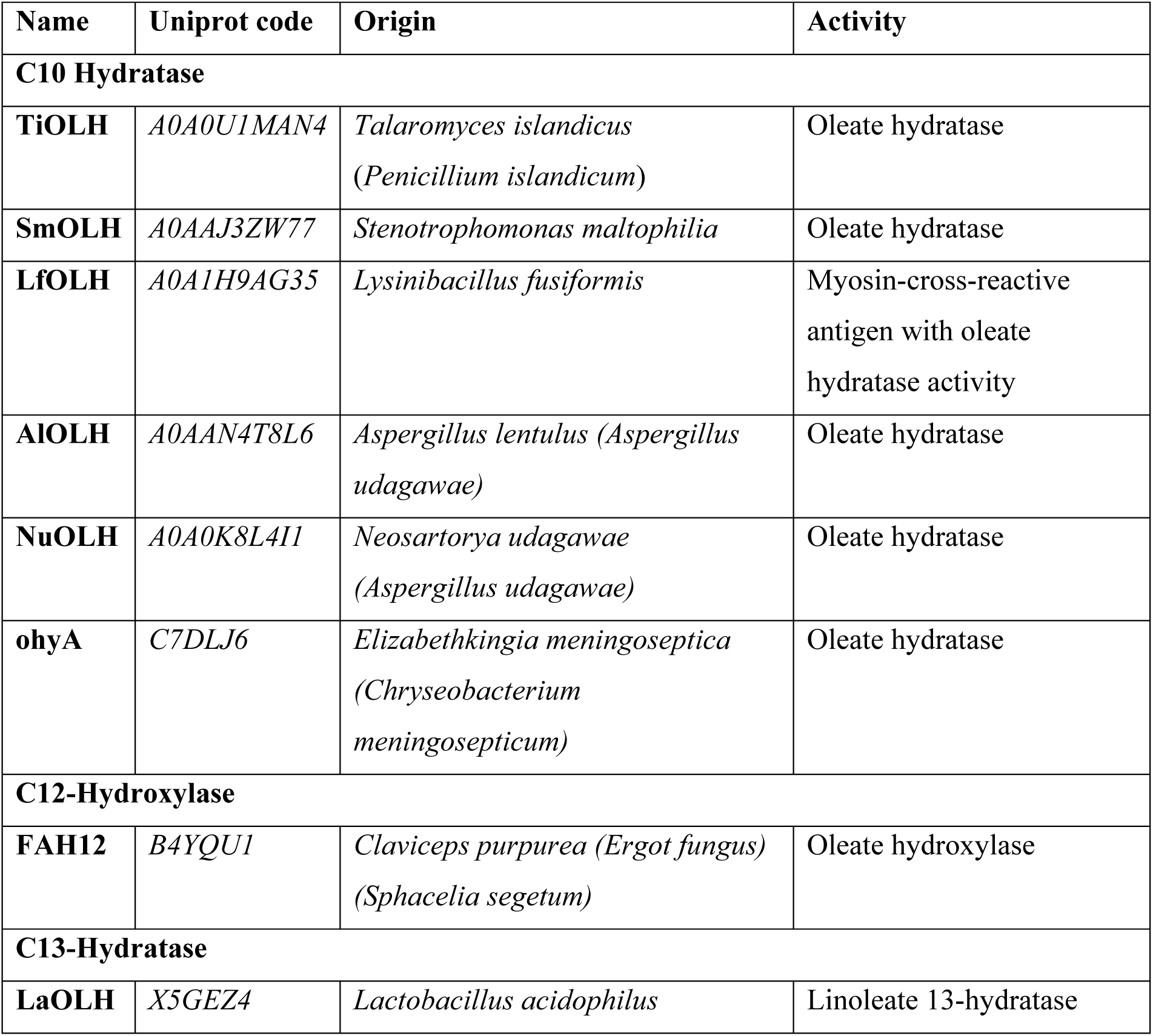
List of enzymes used for hydroxylating fatty acids in yeast.

Since γ-decalactone is the focus of this study, CpFAH12 was selected for further characterization. To confirm the ability of *S. cerevisiae* expressing CpFAH12 from *GAL1* promoter to produce ricinoleic acid, we cultivated the *pox1Δ PGAL-CpFAH12* strain for 24 h in YEP-gal medium supplemented with 1 mM oleic acid and samples were analyzed for both intracellular and extracellular ricinoleic acid. *POX1* (acyl-CoA oxidase gene) was deleted to prevent degradation of ricinoleic acid to lactones for quantitative estimation of ricinoleic acid. The externally supplied 1 mM oleic acid was consumed within 24 h to produce ∼2.5 µmole ricinoleic acid, the majority of which was found to be extracellular (Figure 2).

**Figure 2.**
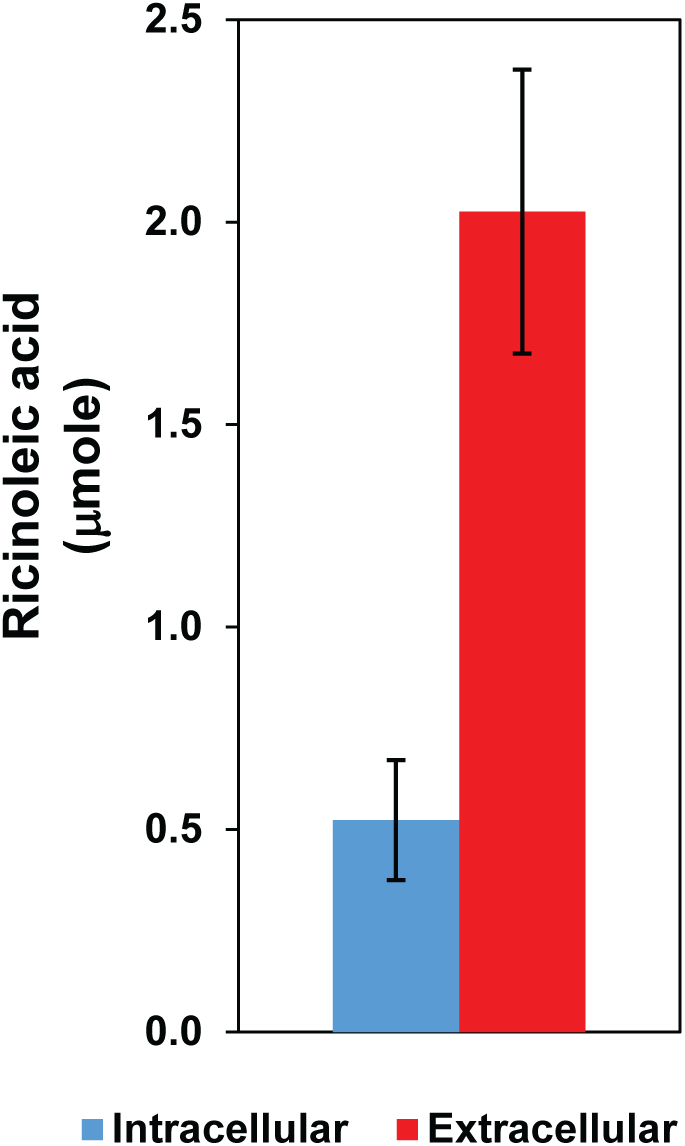
Ricinoleic acid produced in yeast cells expressing CpFAH12 is mainly extracellular The *pox1Δ GAL-CpFAH12* strain was grown for 24h in YEP-gal medium supplemented with 1mM oleic acid and samples were analyzed for both intracellular and extracellular ricinoleic acid.

### Optimization of cultivation parameters for γ-decalactone synthesis in *S. cerevisiae*

For efficient production of γ-decalactone, the extent and the timing of CpFAH12 expression with regards to the growth cycle is critical. Constitutively active promoters have been studied extensively in the context of alcohol and fatty acid synthesis or the derivatives thereof (14–17). The 571bp *ADH2* promoter of *S. cerevisiae* is constitutive, shows higher expression compared to *CUP1*, *GAL1* or *PGK1* promoters and has been effectively used in synthesis of polyketides and triacetic acid lactone (18, 19). *ADH2* is also a glucose-repressible promoter; it is active when glucose in the growth medium is exhausted and ethanol consumption commences (20, 21). The use of *ADH2* promoter to drive CpFAH12 expression provides two distinct advantages for lactone production: (i) separation of growth and lactone synthesis phases which would allow full biomass accumulation before *PADH2-CpFAH12* construct is expressed to induce lactone synthesis (ii) avoidance of lactones’ inhibitory effect on growth as lactone production commences only after cells reach stationary phase. To test the efficiency and the timing of lactone production under these conditions, a strain harboring *PADH2-CpFAH12* was cultivated in 1% (w/v) or 2% (w/v) glucose-supplemented YEP medium containing 1mM oleic acid and lactone accumulation was monitored at 24, 48 and 72 h. In 1% glucose, while no γ-decalactone was detected at 24 h, the maximal yield (∼12 mg/L) was seen at 48 h which remained at this level after 72 h. In 2% glucose, γ-decalactone was detected at low levels at 48 h but increased to ∼10 mg/L at 72 h (Figure 3A). This is consistent with the notion that high glucose content in the medium takes longer to be fully utilised by cells before *ADH2* promoter is de-repressed. In subsequent experiments, we used *ADH2* promoter driven CpFAH12 expression unless mentioned otherwise.

**Figure 3.**
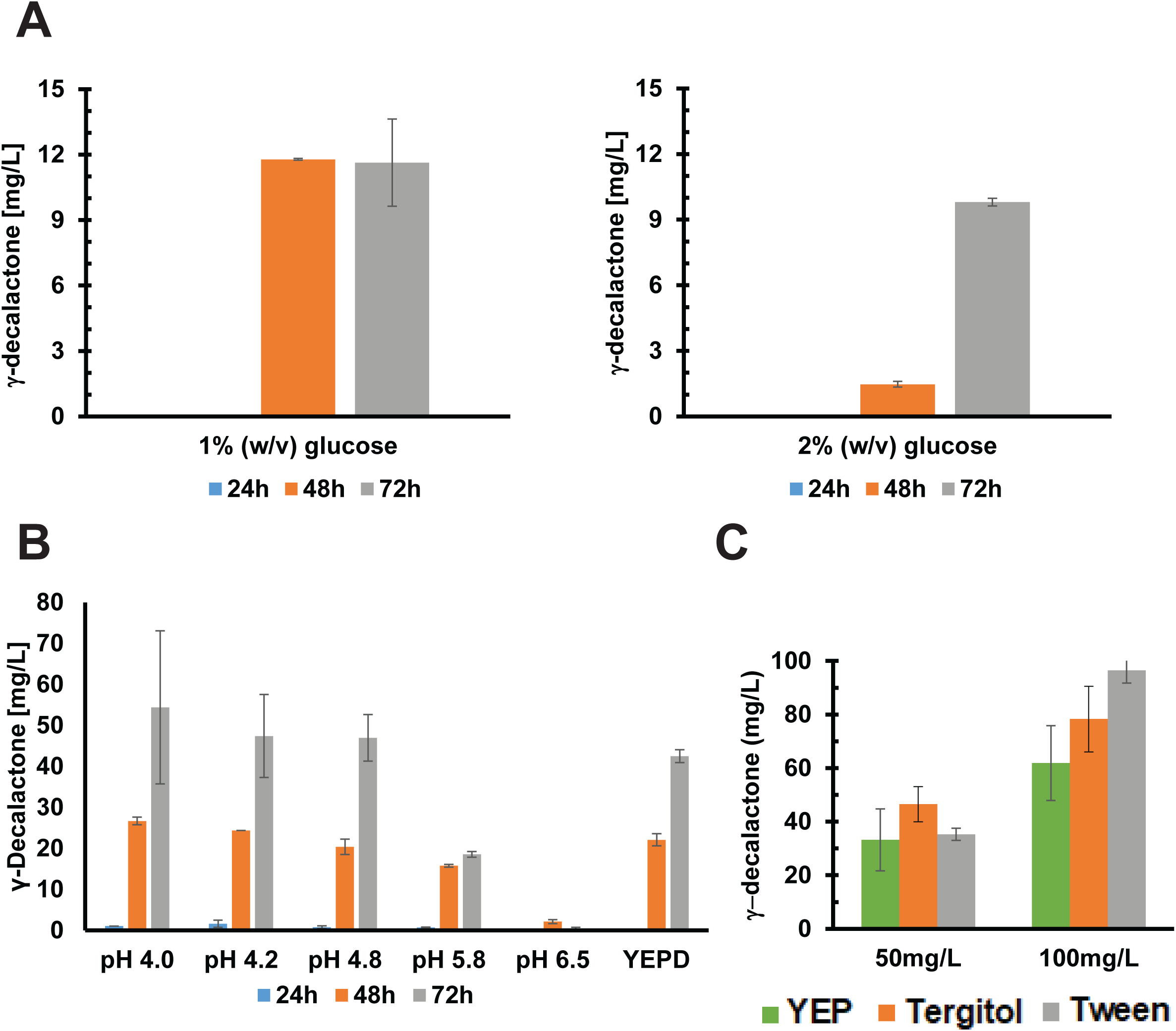
Optimizing the culture conditions and media for the production of γ-decalactone A) Synthesis profile of γ-decalactone with *PADH2-CpFAH12* in *S. cerevisiae* cultured in the presence of 1%(w/v) glucose or 2%(w/v) glucose. The experiment was performed with biological and technical duplicates. B) Effect of pH control on lactone production in *S. cerevisiae PADH2*-*CpFAH12*. The YEPD medium was adjusted with citric acid and dipotassium phosphate to pH 4, pH 4.2, pH 4.8, pH 5.8 and pH 6.5. C) Remaining γ-decalactone in YEP medium after 72h at 24°C, when Tween or Tergitol were supplemented. The initial concentrations of γ-decalactone were 50mg/L and 100mg/L

Since cell viability should be optimum for the efficiency of biotransformation process, we also sought to determine the optimum pH for γ-decalactone production. The hydroxy fatty acid exists in a protonated form in the cultivation broth like YEPD and could interact with membrane lipids. The levels of the protonated form of 4-hydroxydecanoic acid increases with decreasing culture pH and may potentially result in membrane leakage and reduced growth rate (22). Spontaneous lactonization of 4-hydroxydecanoic acid is also problematic and can lead to unwanted effects. Therefore, we tested a range of buffered YEP media (pH 4-6.5) supplemented with 1% glucose to determine the most suitable pH for lactone synthesis. In all cases, adjusting the pH below 4.8 at the start of cultivation leads to lactone production without significantly affecting biomass accumulation (Figure 3B and data not shown). Using medium buffered to pH 5.8 or 6.5 was not beneficial as it caused reduced cell growth. Therefore, growth medium of pH 4.75 was used for all subsequent experiments.

γ-decalactone is a semi-volatile compound with relatively low vapor pressure (0.0±0.5 mm Hg at 25 °C), low Henry’s law constant (3.933 x 10^-6^atm-m^3^/mole) and high boiling point (266 °C). To reduce the loss of lactone (up to 35%) during its synthesis, Tween80 and Tergitol NP-40 were tested for their ability to stem the loss of lactone due to volatility. Tween80 has been used in biotransformation experiments with *Y. lipolytica* to improve the solubility and increase uptake of ricinoleic acid (23). YEP medium (without any cells) with or without 50 mg/L or 100 mg/L of Tergitol or Tween80 was incubated at 24 °C for 72 h. The loss of γ-decalactone was found to be lowest in YEP medium containing Tween80 (Figure 3C). Therefore, Tween80 (100 mg/L) was used in all subsequent experiments conducted for the determination of lactone yields.

### Genes involved in tryptophan biosynthesis and transport provide resistance to γ-decalactone -mediated toxicity in yeast cells

Biotransformation processes can be limited by product-induced toxicity to the host organism. Lactones have been described to have antimicrobial properties because of their ability to interact with cell membranes due to their hydrophobic nature (24). γ-decalactone is detrimental to growth of *Y. lipolytica* at concentrations higher than 150 mg/L (24). Hence, we wished to address and mitigate these factors before investigating metabolic barriers to oleic acid-to-lactone conversion. To determine antimicrobial effects of γ-decalactone and γ-dodecalactone on *S. cerevisiae*, the minimum inhibitory concentration (MIC) was determined. While ∼83% cells survived at 625 µM (106 mg/L) γ-decalactone, less than 10% of cells survived in the presence of 1250 µM (212 mg/L) (Figure 4A). Among the hydrophobic lactones, γ-dodecalactone is known to have a strong inhibitory effect on cell growth (24). Consistent with this, only 32% cells survived in the presence of 156 µM (30.9 mg/L) γ-dodecalactone, while the inhibition was >90% in the medium containing 312 µM (61.8 mg/L) (Figure 4B).

**Figure 4.**
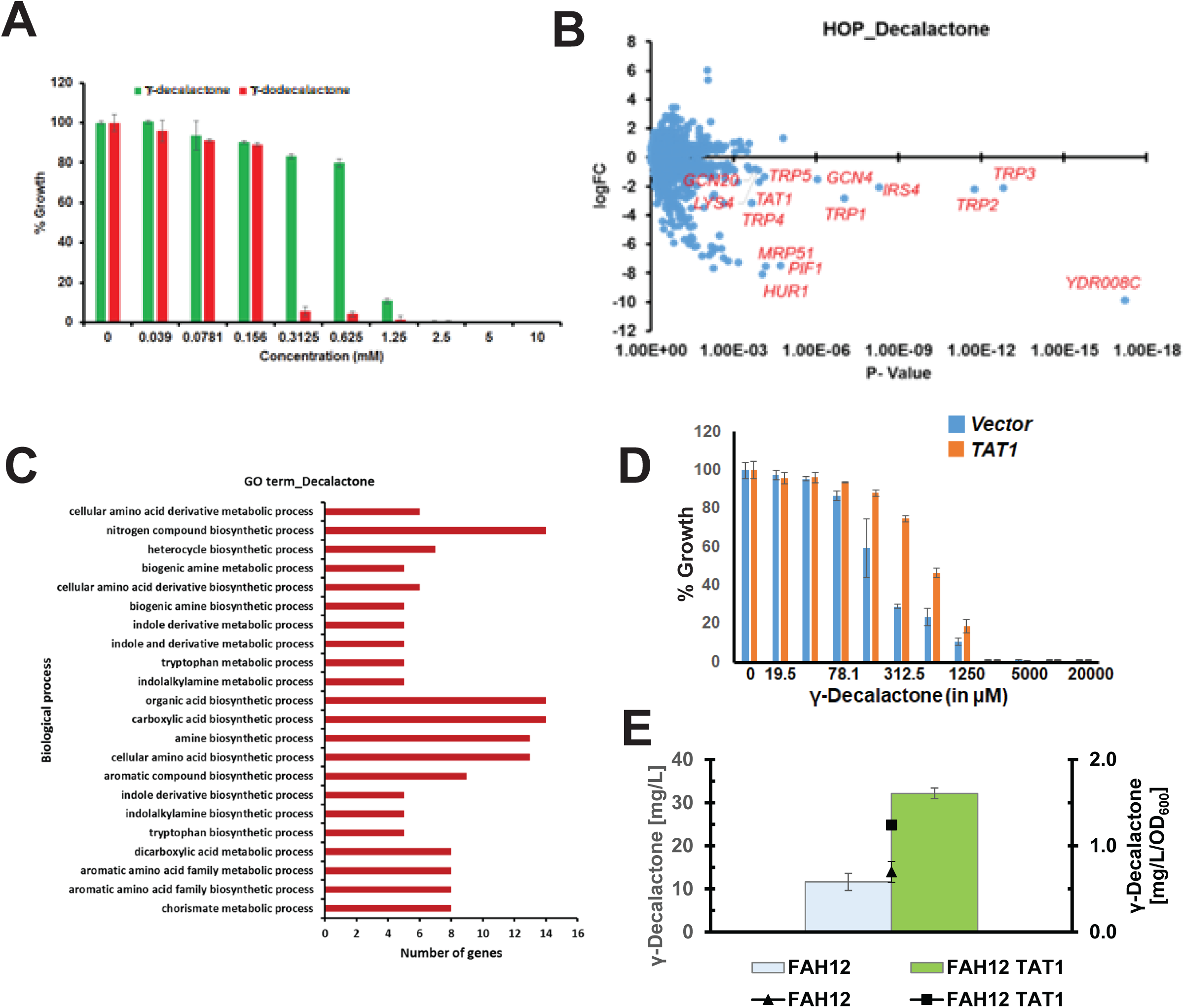
Chemogenomic profiling of γ-decalactone in yeast A) Growth of wild type yeast BY4743 was assayed in the presence of varying concentrations of γ-decalactone and γ-dodecalactone in rich medium (YPD). B) Plot of logarithm of fitness coefficient (logFC) versus P-value derived from homozygous profiling (HOP) assay performed with γ-decalactone. 101 mutants were obtained with logFC < − 0.5 and P-value < 0.05. Gene names of a few mutants are indicated in the plot. C) Results of Gene Ontology enrichment analysis of the 101 hit genes performed with DAVID is depicted. D) γ-Decalactone sensitivity of yeast cells either overexpressing Tat1 or containing an empty vector E) Effect of expressing the amino acid transporter Tat1 on γ-decalactone yields and growth. Increase in titer is mostly due to better cell growth.

To identify genes involved in the modulation of lactone-induced toxicity, we performed chemogenomic profiling in *S. cerevisiae* which exploits the bar-coded nature of the ‘yeast knockout (KO) to determine the targets of bioactive compounds (25–30). We used the pooled yeast haploid knockout strain collection that contains ∼5000 yeast strains, each bearing a deletion of a non-essential gene with a unique barcode. We incubated the pooled yeast gene deletion library in YPD medium in the presence of either γ-decalactone (200 µM) or γ-dodecalactone (50 µM) or DMSO and allowed the cells to grow for five generations. We performed Next Generation Sequencing of the barcodes flanking the gene deletions from the genomic DNA prepared from the three cultures to identify mutants that are sensitive to γ-decalactone and γ-dodecalactone. About 101 mutants were found to be sensitive to γ-decalactone as defined by log FC (Fitness Coefficient) < -0.5 and P-value < 0.05. To gain insights into inhibitory effect of lactones, we performed Gene Ontology analyses of the 101 resistance genes that increase sensitivity to γ-decalactone using Database for Annotation, Visualization and Integrated Discovery (DAVID) (31, 32). Biological process enrichment analysis revealed that several γ-decalactone resistance gene products were involved in tryptophan biosynthesis or transport (Figure 4C). Similar results were obtained with γ-dodecalactone (Supplementary Figure S1). *TAT1* encodes an amino acid transporter that exhibits high affinity tyrosine but low affinity for tryptophan (33). Among the various hits, we found that overexpression of Tat1 in BY4743 strain boosted resistance to γ-decalactone (Figure 4D and data not shown). To test whether overexpression of Tat1 also influences lactone yield, a strain carrying *PADH2-CpFAH12* at *YRPCΔ15* locus with or without *TEF1-*driven *TAT1* gene at its native locus was grown in the presence in YEPD medium containing 1 mM oleic acid and content of extracellular lactone was determined after 72 h. An increase of 63% in γ-decalactone yield was observed (Figure 4E). Cells expressing Tat1 also showed improved growth compared to the control. When corrected for improved growth, the increase in γ-decalactone yield was ∼43%. Thus, about 31% of the improvement in yield was due to enhanced growth due to Tat1 overexpression. It is possible that improved lactone yield by Tat1 overexpression is due to both enhanced growth and mitigation of lactone toxicity. However, the mechanism by which Tat1 dampens lactone toxicity is not clear at present.

### Impact of various metabolic pathway branches and effectors on overall synthesis of γ- decalactone

We then aimed to delineate the influence of various metabolic branches on γ-decalactone synthesised from the conversion of de novo pool of oleic acid and not from externally supplemented oleic acid. This approach excludes the complications related to oleic acid transport and uptake. The intracellular synthesis of fatty acids requires multiple steps involving the fatty acid synthase complex subunits (Fas1 and Fas2) and the acetyl-CoA carboxylase (Acc1). The precursor of acetyl-CoA (pyruvate) is also utilised in other metabolic pathways that compete with the fatty acid biosynthesis. The formation of NADPH, as a reduction equivalent, is also essential for activity of the fatty acid synthase complex. Since acyl-CoA can be stored in form of triacylglycerides or sterol ester until the carbon source is depleted, the storage of fatty acids also represents an additional competing pathway. Figure 5A depicts the main pathways and side products as gleaned from the existing literature.

**Figure 5.**
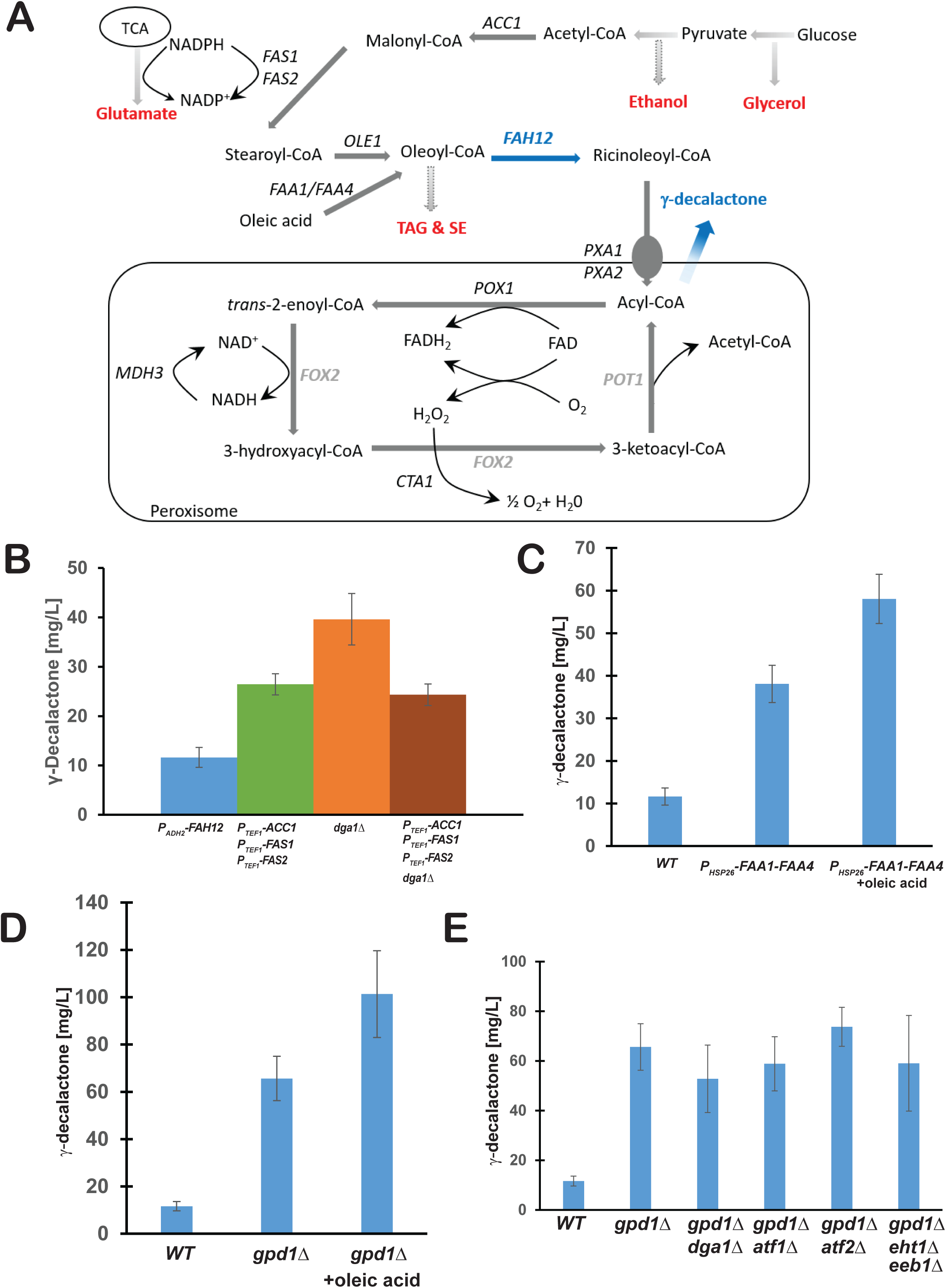
Boosting the metabolic flux to enhance γ-decalactone production in yeast A) Metabolic pathways identified for optimizing γ-decalactone precursors in *S. cerevisiae* from the literature. *ACC1* (acetyl-CoA carboxylase); *FAS1* (fatty acid synthase subunit β); *FAS2* (fatty acid synthase subunit α); *DGA1* (diacylglycerol acyltransferase); *OLE1* (Delta(9) fatty acid desaturase); *POT1*(peroxisomal Oxoacyl Thiolase); *FAA1-4* (Fatty acyl CoA synthetase). *POX1* (Fatty-acyl coenzyme A oxidase); *FOX2* (3-hydroxyacyl-CoA dehydrogenase and enoyl-CoA hydratase); *CTA1* (Catalase A); *PXA1/2* (Peroxisomal ABC-transporter); TAG (Triacylglycerol); SE (Steryl ester). B) Effect of either overexpressing *ACC1* (acetyl-CoA carboxylase), *FAS1* (fatty acid synthase subunit β), and *FAS2* (fatty acid synthase subunit α) or deleting *DGA1* (diacylglycerol acyltransferase) or both of the above manipulations on γ-decalactone production was assayed. All strains are expressing *CpFAH12* with the *ADH2* promoter in YEPD + 1% w/v Glucose. pH was adjusted to 4.75 using 100mM citric acid and 100mM K2HPO4 C) Overexpression of long chain fatty acyl-CoA synthetases Faa1 and Faa4 improved γ- decalactone titers with and without supplementation of 1mM oleic acid. D) γ-Decalactone synthesis in *S. cerevisiae* mutants deficient in glycerol formation (*GPD1*), E) Effect of deleting *GPD1* in combination with deletions in *DGA1* (Diacylglycerol acyltransferase), or *ATF1* (Alcohol acetyltransferase) or *ATF2* (Alcohol acetyltransferase), or *EHT1* (Acyl-coenzymeA:ethanol O-acyltransferase) + *EEB1* (Acyl-coenzymeA:ethanol O-acyltransferase) on γ-decalactone production.

Augmenting the fatty acid synthesis is an obvious choice for providing greater amount of the precursor oleic acid for CpFAH12 to produce 12-hydroxy-cis-9-octadecenoic acid (ricinoleic acid). It has been reported that the overexpression of all three fatty acid biosynthesis genes (*FAS1, FAS2, ACC1*) lead to 70.6 mg/L total fatty acid content and an increase in more than 39% compared to the wild type strain (15). To increase the availability of substrate for CpFAH12, we expressed native *FAS1*, *FAS2* and *ACC1* from the *TEF1* promoter (by promoter swapping) in a strain harboring *PADH2-CpFAH12* and measured the extent of γ-decalactone production. The effect was significant with a 2.3-fold increase in overall yield (from 11.6 to 26.4 mg/L) of γ-decalactone compared to the control strain (Figure 5B). This suggests that the availability of the precursor oleic acid is limiting. Since triglyceride synthesis is an important metabolic diversion of intracellular fatty acids (Figure 5A), we also analyzed the effect of the deletion of *DGA1* gene encoding diacylglycerol acyl transferase, the enzyme that catalyzes the acylation of diacylglycerol. The deficiency of *DGA1* showed a 3.4-fold increase (from 11.63 to 39.6 mg/L) in γ-decalactone production compared to the control strain (Figure 5B). To test if the combination of *DGA1* deficiency with overexpression of *FAS1*, *FAS2* and *ACC1* would enhance the metabolic flux towards ricinoleic acid synthesis and consequently, the synthesis of γ-decalactone, we deleted *DGA1* gene in the strain overexpressing *FAS1*, *FAS2* and *ACC1* from *TEF1* promoter. Interestingly, combination of these elements failed to enhance γ-decalactone synthesis which remained at a level (24.3 mg/L) comparable to that in *FAS1*, *FAS2* and *ACC1* overexpressing strain (Figure 5B).

Next, we examined the effect of genes involved in the import and activation of exogenously supplied fatty acids on γ-decalactone synthesis. *S. cerevisiae* can survive on medium- and long-chain fatty acids as the sole carbon source by importing the fatty acids and subsequently catabolizing them via β-oxidation in the peroxisomes (34). The fatty acyl-CoA synthetases have a role in competitive import of long chain fatty acids and their activation (34). Although five fatty acyl-CoA synthetases have been described (Faa1-4 and Fat1), most of the activity towards exogenous fatty acids results from Faa1 and Faa4 (35, 36). Thus, we overexpressed *FAA1* and *FAA4* under the control of *HSP26* promoter in a strain harboring *PADH2-CpFAH12* and cultured for 72 h with and without supplementation of oleic acid. The γ-decalactone yield increased significantly by overexpression of *FAA1* and *FAA4* (from 38.1 mg/L and 58.1mg/L) (Figure 5C). It is interesting to note that even in the absence of exogenous oleic acid, overexpression of *FAA1* and *FAA4* results in higher yield of γ-decalactone (38.1 mg/L) compared to the control strain i.e. without overexpression of these genes (11.63 mg/L; Fig 5C). It is possible that *FAA1/FAA4* overexpression increases the de novo synthesised oleic acid flux towards ricinoleic acid. However, currently there is no clear metabolic rationale for this notion.

### Glycerol-3-phosphate dehydrogenase-mediated glycerol synthesis pathway dampens γ- decalactone production

The intracellular pool of acetyl-CoA is important for the de novo fatty acid synthesis. Improvement of acetyl-CoA availability by interrupting ethanol or glycerol synthesis has been shown to enhance the production of fatty alcohols, alkanes or fatty aldehydes (18, 37–39). Glycerol production, though important for adaptation to osmotic stress, is considered energetically wasteful during normal growth conditions in *S. cerevisiae*, since it functions to dispose excess reducing power (40). The NAD^+^-dependent glycerol 3-phosphate dehydrogenase (GPD) catalyzes the reduction of dihydroxyacetone phosphate to glycerol 3-phosphate and thus constitutes the first step in glycerol synthesis (40, 41). In *S. cerevisiae*, NAD^+^-dependent glycerol 3-phosphate dehydrogenase is encoded by two homologous genes *GPD1* and *GPD2* (41). Deletion of *GPD1* gene leads to significant reduction in glycerol synthesis (42). Therefore, we asked if *GPD1* deficiency would not only enhance intracellular pool of acetyl-CoA and consequently, lactone production, but also augment the conversion of externally supplied oleic acid to γ-decalactone by stemming the diversion of oleic acid to triacylglycerol synthesis. *PADH2-CpFAH12 gpd1Δ* strain was grown in YEPD medium (1% w/v glucose) with or without exogenous oleic acid and γ- decalactone production was measured after 72 h. The lactone yield in *PADH2-CpFAH12 gpd1Δ* was 5.6-fold higher (at 65.2 mg/L) compared to the *PADH2-CpFAH12* strain (11.6 mg/L) (Figure 5D). Cultivation in the presence of exogenous oleic acid (1mM) further increased the lactone yield to ∼100 mg/L). Thus, glycerol synthesis pathway is a significant metabolic diversion that negatively impacts the lactone yield.

*S. cerevisiae* is used in wine and beer production, because of the high activity of alcohol and acyl-CoA acetyltransferase. These pathways consume ethanol and acetate, thus acetyl-CoA. Therefore, we tested the effect of deficiencies of the following genes involved in these pathways in combination with *gpd1*Δ: *DGA1* (Diacylglycerol acyltransferase), *ATF1*/*ATF2* (Alcohol acetyltransferases), *EHT1* (Octanoyl-CoA:ethanol O-acyltransferase), *EEB1* (Acyl-coenzymeA:ethanol O-acyltransferase). *gpd1Δ dga1Δ*, *gpd1Δ atf1Δ*, *gpd1Δ atf2Δ*, *gpd1Δ eht1Δ eeb1Δ* strains were cultivated in YEPD (1% glucose w/v) without exogenous oleic acid for 72 h. Except *gpd1Δ atf2Δ*, none of the other combinations surpassed the lactone yield exhibited by *gpd1Δ* strain (Figure 5E). Though the lactone yield in *gpd1Δ atf2Δ* was higher than in *gpd1Δ* strain, the increase was marginal.

### Fatty acyl synthetase Faa2 is essential for biotransformation of oleic acid to γ-decalactone

The intracellular transport of oleic acid is an important element in the biotransformation of exogenous oleic acid to γ-decalactone. Fatty acid transport is generally thought to be coupled to its intracellular activation via esterification to coenzyme A by fatty acyl-CoA synthetases. *S. cerevisiae* express five fatty acyl-CoA synthetases encoded by *FAA1*, *FAA2*, *FAA3*, *FAA4* and *FAT1* genes (43). Cells deficient in each of the five genes are viable. While Faa1/Faa4 are cytoplasmic and are involved in the activation of long chain fatty acids, Faa2 is localised in the peroxisomes and is involved in the activation of medium chain fatty acids for peroxisomal β- oxidation (44). Fat1 is required for activation of very long chain fatty acids (36). The specific role of Faa3 is thus far unclear.

To investigate the role of fatty acyl synthetases, we constructed *gpd1Δ PADH2-CpFAH12 faa1Δ*, *gpd1Δ PADH2-CpFAH12 faa2Δ*, *gpd1Δ PADH2-CpFAH12 faa4Δ* and *gpd1Δ PADH2-CpFAH12 faa1Δ faa4Δ* strains. The strains were cultivated in YEPD medium supplemented with 1 mM oleic acid for 72 h at 24 °C and γ-decalactone content in the medium was determined by GC-MS. The *gpd1Δ PADH2-CpFAH12* strain served as a reference. As seen in our previous experiments, the γ- decalactone yield in *gpd1Δ PADH2-CpFAH12* strain was ∼90 mg/L (Figure 6A). The deficiency of Faa1 or Faa4 reduced the γ-decalactone yield moderately to ∼60 mg/L and ∼78 mg/L, respectively. Deleting both Faa1 and Faa4 reduced the γ-decalactone yields to 22 mg/L. Surprisingly, the deficiency of Faa2 completely abolished the conversion of oleic acid to γ-decalactone (Figure 6A). We also examined the levels of oleic acid and ricinoleic acid in the strains. Oleic acid level in the *faa1Δ faa4Δ* culture supernatant was 20-fold higher than in the wild type culture supernatant indicating that Faa1 and Faa4 are required for efficient utilization of oleic acid. However, ricinoleic acid levels in the *faa1Δ faa4Δ* strain were not very different from that of the wild type strain. In contrast to the *faa1Δ faa4Δ* strain, levels of ricinoleic acid but not oleic acid accumulated in the *faa2Δ* strain indicating that in the absence of Faa2, ricinoleic acid cannot be converted to γ- decalactone. These results indicate that Faa2 is essential for γ-decalactone production in yeast.

**Figure 6.**
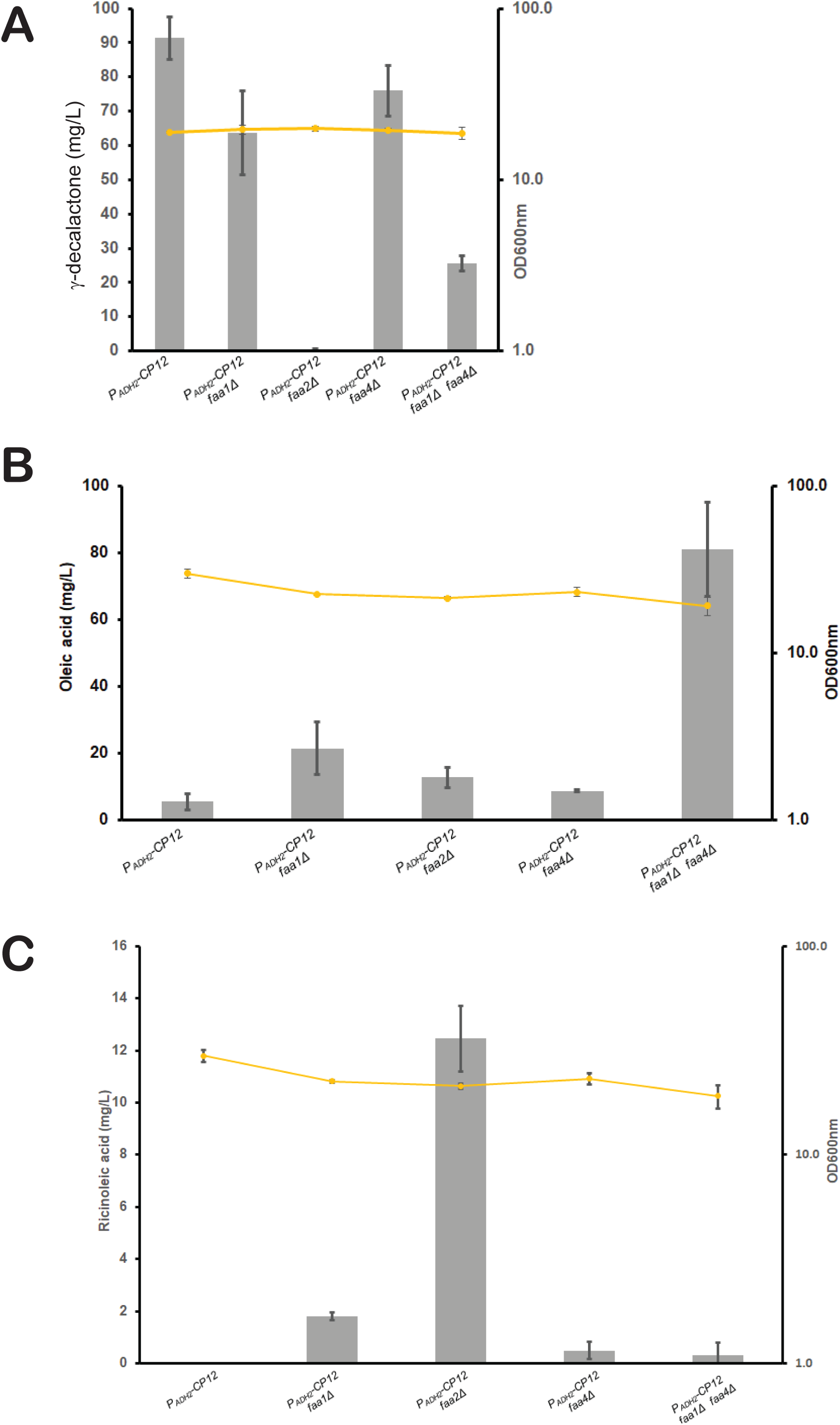
Faa2 is essential for γ-decalactone production from ricinoleic acid in yeast Yeast cells overexpressing CpFAH12 and containing either wild type or *faa1Δ*, *faa4Δ*, *faa2Δ* or *faa2Δ faa4Δ* were grown in buffered YEPD medium + 1% w/v glucose and oleic acid. Levels of γ-decalactone (A), oleic acid (B) and ricinoleic acid (C) measured by GC-MS are plotted.

### Pxa1/2 and peroxisomal localization of Faa2 are required for efficient γ-decalactone production in yeast

Our observations are consistent with the hypothesis that Faa2 is required for the conversion of ricinoleic acid to ricinoleoyl-CoA which then undergoes β-oxidation to produce γ-decalactone in the peroxisomes. This may appear somewhat surprising as the cytosolic Faa1 and Faa4 would be expected to convert ricinoleic acid to ricinoleoyl-CoA. Interestingly, it has been proposed that fatty acyl CoA transported to the peroxisomes via the ABC transporter proteins Pxa1/Pxa2 are hydrolyzed and esterified back into the CoA form in the peroxisomes prior to β-oxidation (45). If this notion were correct, then either deleting *PXA1/PXA2* or blocking the peroxisomal localization of Faa2 should severely affect γ-decalactone production. We compared the levels of γ-decalactone, oleic acid and ricinoleic acid levels in wild type, *pxa1Δ*, *pxa2Δ* and *pxa1Δ pxa2Δ* strains expressing CpFAH12. Consistent with our prediction, the γ-decalactone levels were reduced by 2-2.3-fold in the *pxa1Δ*, *pxa2Δ* and *pxa1Δ pxa2Δ* strains compared to the wild type strain (Figure 7A). Ricinoleic acid levels in the *pxa2Δ* and *pxa1Δ pxa2Δ* strains were slightly elevated compared to the wild type and *pxa1Δ* strains. To test whether the peroxisomal localization of Faa2 is required for γ- decalactone production from ricinoleic acid in yeast, we first identified the Peroxisomal Targeting Sequence (PTS) of Faa2 using the online protein localization sequence predictor tool DeepLoc 2.0 (46). The four amino acid residue motif TEKL at the C-terminus of Faa2 was identified as the peroxisomal targeting sequence with a probability of 0.9773 (Supplementary Figure S2). Deletion of the c-terminal peroxisomal targeting sequence affects peroxisomal localization of proteins in yeast (47, 48). We therefore deleted the entire TEKL motif or replaced lysine in the motif with arginine. As expected, while the wild type Faa2 strain produced γ-decalactone, the *faa2Δ* mutant strain failed to do so. Strains expressing the Faa2 mutants with defective peroxisomal targeting sequence failed to produce γ-decalactone and accumulated ricinoleic acid (Figure 7B). Thes results strongly suggest that the peroxisomal localization of Faa2 is essential for γ-decalactone production from ricinoleic acid in yeast.

**Figure 7.**
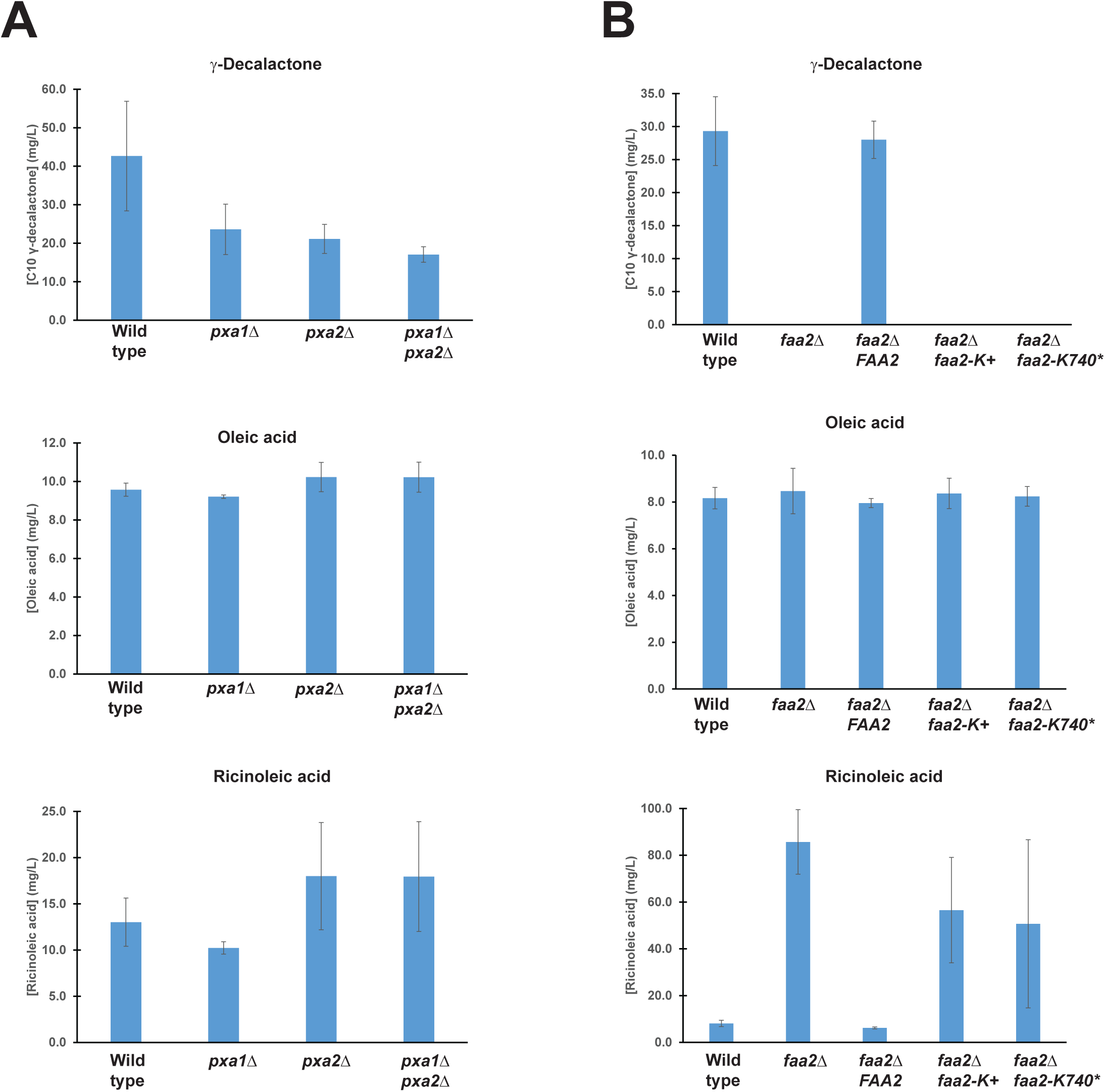
Pxa1/2 and peroxisomal localization of Faa2 is essential for efficient γ- decalactone production from ricinoleic acid in yeast A) Yeast cells overexpressing CpFAH12 and containing either wild type *PXA1 PXA2* or *pxa1Δ* or *pxa2Δ* or *pxa1Δ pxa2Δ* mutants were grown in buffered YEPD medium + 1% w/v glucose and oleic acid. Levels of γ-decalactone, oleic acid and ricinoleic acid in the different strains measured by GC-MS are plotted. B) Yeast cells overexpressing CpFAH12 and containing either wild type or *faa2Δ* or *faa2Δ* mutants were grown in buffered YEPD medium + 1% w/v glucose and oleic acid. Levels of γ-decalactone, oleic acid and ricinoleic acid in the different strains measured by GC- MS are plotted.

## Discussion

Driven by the need to establish a non-toxic strategy to produce γ-decalactone, we combined metabolic engineering and chemogenomic profiling methods to establish a platform for producing flavour lactones from oleic acid in *S. cerevisiae.* Blocking the conversion of Dihydroxyacetone phosphate (DHAP) to glycerol by deleting *GPD1* had the maximal effect on boosting γ- decalactone yields. Overexpression of the tryptophan transporter Tat1 mitigated lactone toxicity and enhanced lactone production. Finally, our study has uncovered an essential role for the peroxisomal enzyme Faa2 in the conversion of ricinoleic acid to lactones.

We attempted to boost the production of γ-decalactone by diverting the metabolic flux towards oleic acid production. The choice of the glucose-repressible promoter *ADH2* ensured the separation of the biomass growth and lactone production phases. Addition of exogenous oleic acid as a substrate consistently helped to boost lactone production indicating that oleic acid is a bottleneck for lactone production. Multiple lines of additional experimental evidence provide further support for this notion: (i) Overexpression of fatty acid biosynthetic genes *FAS1*, *FAS2* and *ACC1* helped to boost lactone production (ii) reducing the storage of oleic acid in the form of triglycerides by deleting the diacylglycerol acyl transferase gene *DGA1* helped to boost γ- decalactone production by 3.4-fold and (iii) boosting the acetyl CoA levels by limiting the production of glycerol from DHAP by deleting Glycerol-3-phosphate dehydrogenase had a 6-fold effect on γ-decalactone production.

Chemogenomic profiling analyses revealed a surprising link between the tryptophan biosynthesis and uptake pathways and γ-decalactone resistance. Similar results were obtained with the structurally related γ-dodecalactone. It is possible that γ-decalactone binds to the yeast plasma membrane and prevents the uptake of tryptophan by inhibiting the tryptophan transporters, thereby sensitizing the tryptophan-biosynthetic/ transport mutants. Consistent with this idea, overexpression of tryptophan transporter Tat1 enhanced resistance to lactones and increased lactone production. Measuring the uptake of radiolabeled tryptophan in the presence and absence of lactones in yeast should help to test this hypothesis.

The most intriguing finding of our study is the essential role for Faa2 in the conversion of ricinoleic acid to γ-decalactone. Our results are consistent with the hypothesis that ricinoleoyl CoA loses its CoA group during its transportation to peroxisomes which then requires re-acylation by the peroxisomal Faa2 before undergoing β-oxidation. Although it remains to be shown whether deleting the peroxisomal targeting sequence of Faa2 affects its correct localization, similar mutations have been shown to affect the localization of other yeast proteins (47, 48). It was previously reported that the transport of Fatty acyl CoA to the peroxisomes via Pxa1/2 results in the loss of the acyl group (45). Interestingly, they also showed an association between Faa2 and Pxa1/2 (45). Why cells remove and reintroduce the CoA group during the peroxisomal transport of fatty acyl CoA remains enigmatic. It is also possible that free ricinoleic acid is also transported to the peroxisomes via a non Pxa1/Pxa2-dependent mechanism which will require peroxisomal Faa2-mediated acylation for catabolism via β-oxidation. This would also explain the lower levels of γ-decalactone in the *pxa1Δ pxa2Δ* mutant but a complete absence in the *faa2Δ* mutants.

In summary, we have established a yeast-based platform for producing flavour lactones from oleic acid. We strongly believe that our various lactone-boosting strategies can be extended to unconventional yeasts such as *Yarrowia lipolytica* and *Pichia pastoris* and help to generate flavour lactones at a commercial scale.

## Materials and methods

List of strains used in this study and with their genotypes

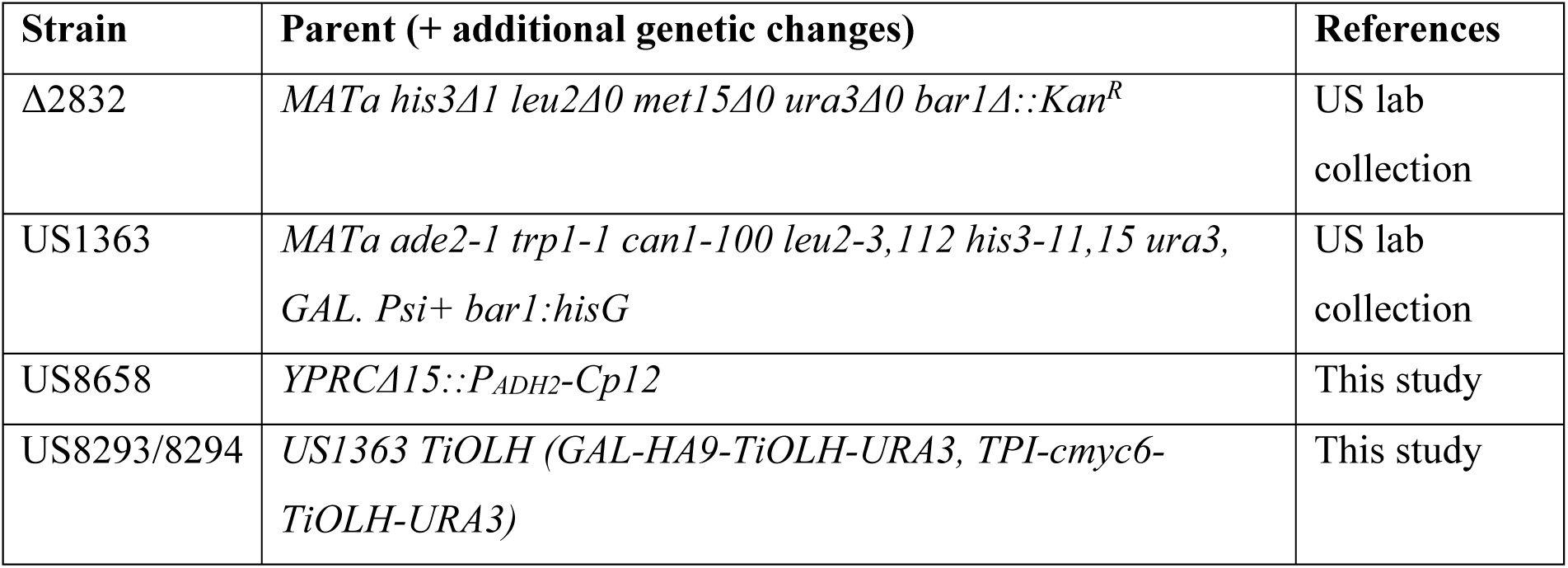
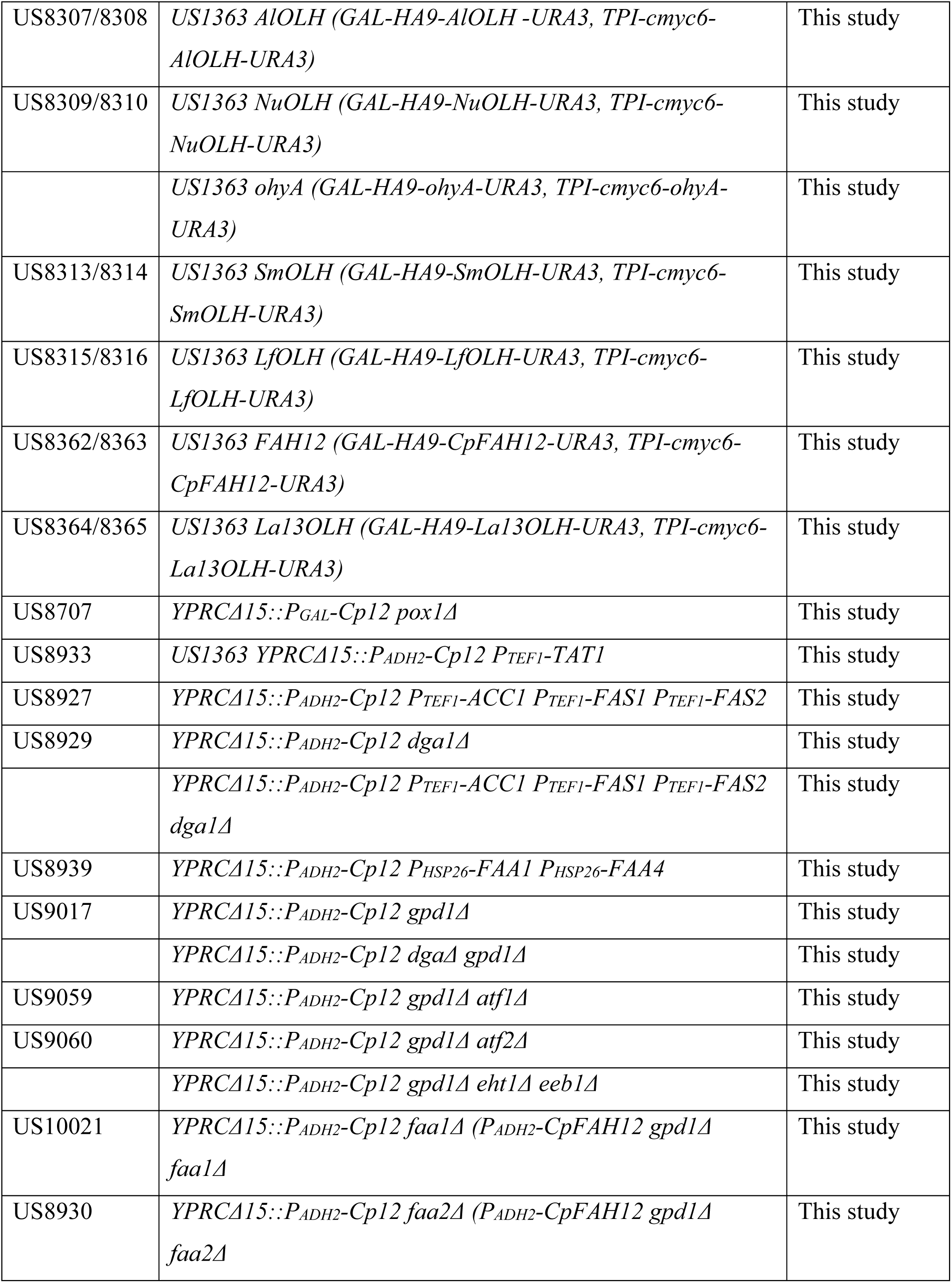
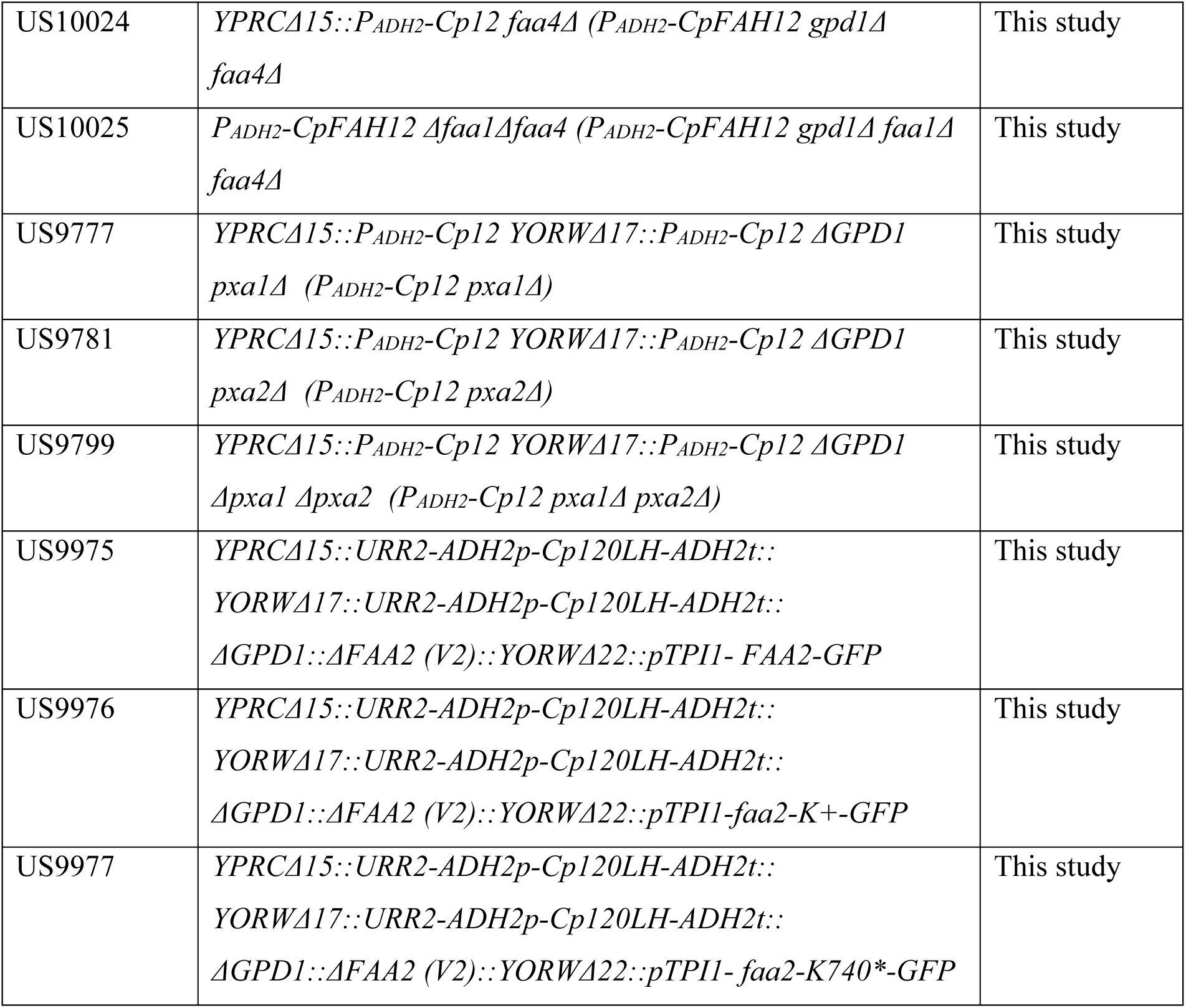

### Strain construction (YeastFab & CrisprCas)

Deletion of genes was performed using the Crispr/Cas9 plasmids according to Generoso et al. 2016 (49). The target sequence was analyzed by the web tool CHOPCHOP (50).

### Media and cultivation conditions

All *E. coli* cells were grown in Gibco LB Broth (ThermoFisher Scientific) with 100 mg/L ampicillin to maintain the corresponding plasmids. *S. cerevisiae* cells were grown in complex (YEP) medium (10 g/l Yeast Extract, 20 g/l Peptone) or synthetic (SC) medium (1.7 g/l yeast nitrogen base (w/o amino acids and ammonium sulphate), 5 g/l ammonium sulphate, amino acids as needed for selection, pH 6.3 (with KOH). For solid medium 20 g/l agar was added before autoclaving. Carbon sources used were added to the medium after autoclaving. Briefly, all *S. cerevisiae* strains were cultivated in 10-20 ml medium for 16 h. The cells were washed with water and used to inoculate a 30 ml culture with a starting optical density (OD600nm) of 0.1 and incubated at 24 °C for another 24-72 h before harvest.

### Minimum Inhibitory Concentration (MIC)

Minimum Inhibitory Concentration (MIC) assessment was performed with the diploid wild type yeast strain BY4743 as described previously (27). Briefly, frozen yeast cells were revived on YPD agar plates and cultured in liquid YPD medium for nine generations (OD600 nm ≤ 2). Cells were diluted to a starting OD600 nm of 0.0625 in YPD medium. 200 μL of the diluted yeast culture was transferred into the 96-well microtiter plate having 2-fold serially diluted concentrations of γ- decalactone/dodecalactone in duplicates. Following incubation of the cultures for 16–24 hours at 30 °C with shaking, the OD600 nm was measured using microplate reader Gen 5^TM^ (BIO-TEK Instrument, Vermont, USA). Minimum inhibitory concentration of lactones was calculated by comparing the growth of treated and untreated cells.

### Chemogenomic profiling

Homozygous profiling assay was performed with the pooled yeast homozygous Knockout collection (Invitrogen) in the BY4743 background, as reported previously (27–29, 51). γ- Decalactone and γ-Dodecalactone were used at 200μM and 50μM respectively which caused a 50% reduction in growth in YPD medium after 16 hours at 30 °C. Preparation of genomic DNA from yeast cells, amplification of barcode sequences by PCR and Next Generation Sequencing (NGS) of the PCR products were all performed as previously described (27, 29).

### Cell Lysate OLH test

Extraction and analysis of lactones and fatty acids (GCs used) For analysis of lactone and fatty acids 250 µl of culture supernatant and an internal standard (2 mg/ml) were vigorously mixed with 250 µl ethyl acetate. After phase separation (3.500 rpm, 5 min), 100 µL organic layer was mixed with BSTFA (N, O-Bistrifluoroacetamide) and incubated for 10 min at 80 °C. The extract was analysed using an Agilent 7890B gas chromatograph equipped with an DB5 capillary column and a mass spectrometer (Agilent 7200 Accurate-Mass Quadrupole Time-of- Flight). A 1 μl portion of sample was injected (split 1:20) at 250°C. The following oven temperature program was applied: 50 °C for 5 min, increase of 10 °C/min to 120 °C and hold for 5 min, increase of 15 °C/min to 180 °C and a hold for 10 min, increase of 20 °C/min to 220 °C followed by a hold for 30 min. Mass spectrometer was operated in EI mode with full scan analysis. Lactones and fatty acids were identified and quantified by comparison with commercially available standards.

### Analysis of fatty acids and lactones from cultures

Fatty acids were methylated before GC-MS analysis. 1ml of culture supernatant was transferred into an eppendorf tube. 1µl of (C17:0) fatty acids was added into the eppendorf tube. 200µl MeOH: Chloroform (1:1) was added to the mixture and the tube was centrifuged at 3500rpm for 5 minutes. About 50µl of the bottom layer was transferred into a 2 mL GC vial and dried in the fume hood overnight. 500µl 1N hydrochloric acid (HCl) in methanol was added to the dried mixture and incubated at 80 ℃ for 1h in an oven. After cooling the samples at room temperature, 500 µl of 0.9% (w/v) NaCl was added to the mixture followed by addition of 500µl of hexane and vortexed for 1 minute. Samples were centrifuged at 3000rpm for 5 minutes and 500µl was taken to a fresh eppendorf tube and centrifuged at 16000rpm for 5 minutes. About 50µl was filtered with Claristep kit into a vial. Lactones were directly extracted from medium using ethyl acetate and analyzed by GC-MS.

### GC-MS

The extracted samples were analysed by GC-MS, performed using Agilent 7890B Gas Chromatograph, coupled with 5977B GC/MSD, equipped with DB-5ms column (Agilent 122- 5532; 30 m x 250 μm x 0.25 μm). The following temperature program was employed 1 μL of sample was injected into the injector set at 250 °C, with a split ratio of 1:10. The helium gas flow was at 1.2 mL/min. The temperature of the column oven was set to the following: an initial temperature of 70 °C, increasing at 20 °C/min to 120 °C, increasing at 10 °C/min to 230 °C, and increasing at 5 °C/min to 280 °C. Samples were ionised with EI in full scan mode from 50-550 Da. Transfer line and ion source were set to 230 °C and 150 °C, respectively, and the collection delay was set to 3 min.

## Acknowledgements

This work was supported by the NRF-NSFC Joint Research Grant (Award number: NRF2018NRF-NSFC003SB-010) awarded to Prof. Uttam Surana. We thank Dr Renata Glitsos and Ms Tiffany Chau for performing a few experiments and for their comments on the manuscript.

## Legends for supplementary figures

**Figure S1.**
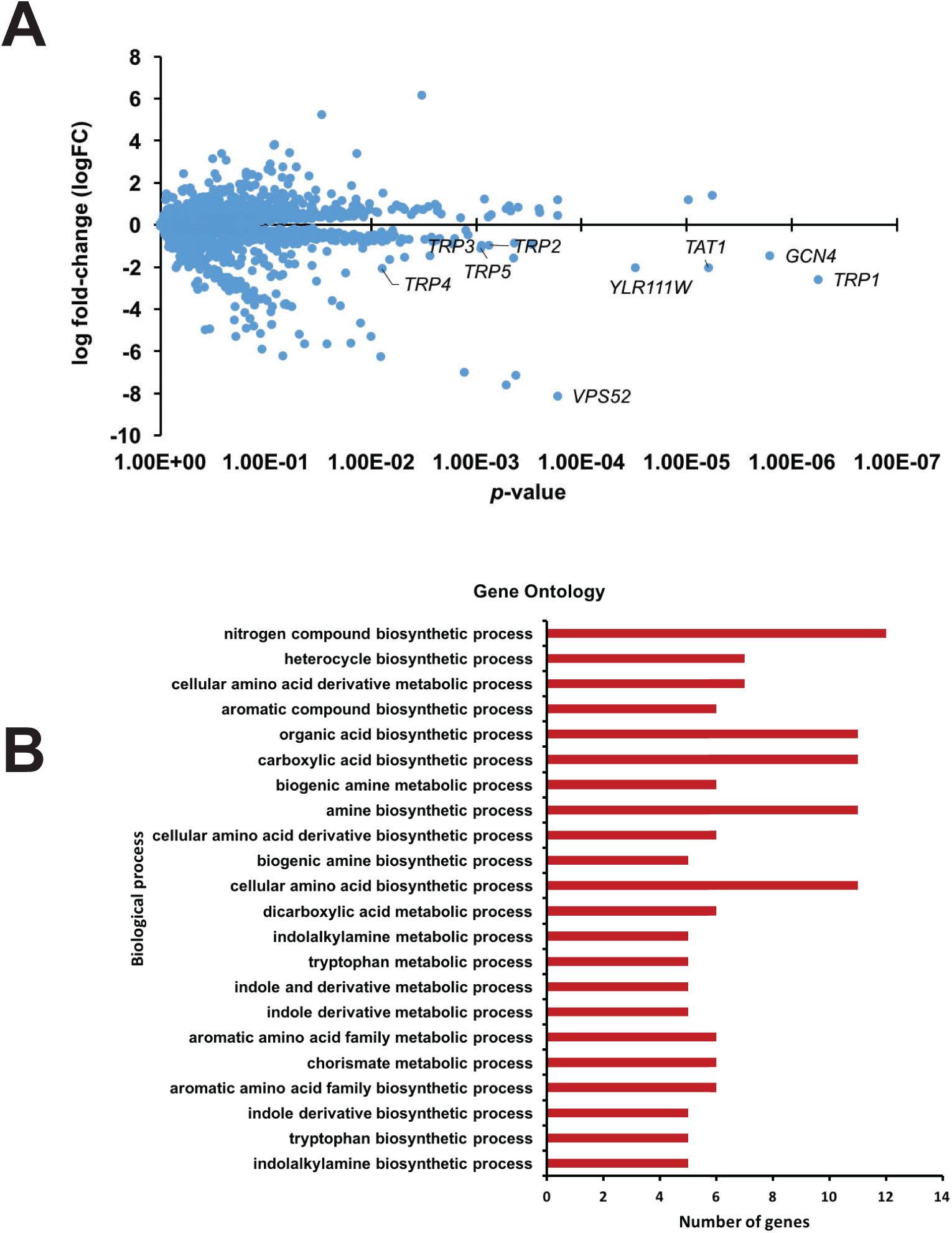
Chemogenomic profiling analysis of γ-dodecalactone A) Plot of logarithm of fitness coefficient (logFC) versus P-value derived from homozygous profiling (HOP) assay performed with γ-dodecalactone. 73 mutants were obtained with logFC < − 0.5 and P-value < 0.05. Gene names of a few mutants are indicated in the plot. B) Results of Gene Ontology enrichment analysis of the 73 hit genes performed with DAVID is depicted.

**Figure S2.**
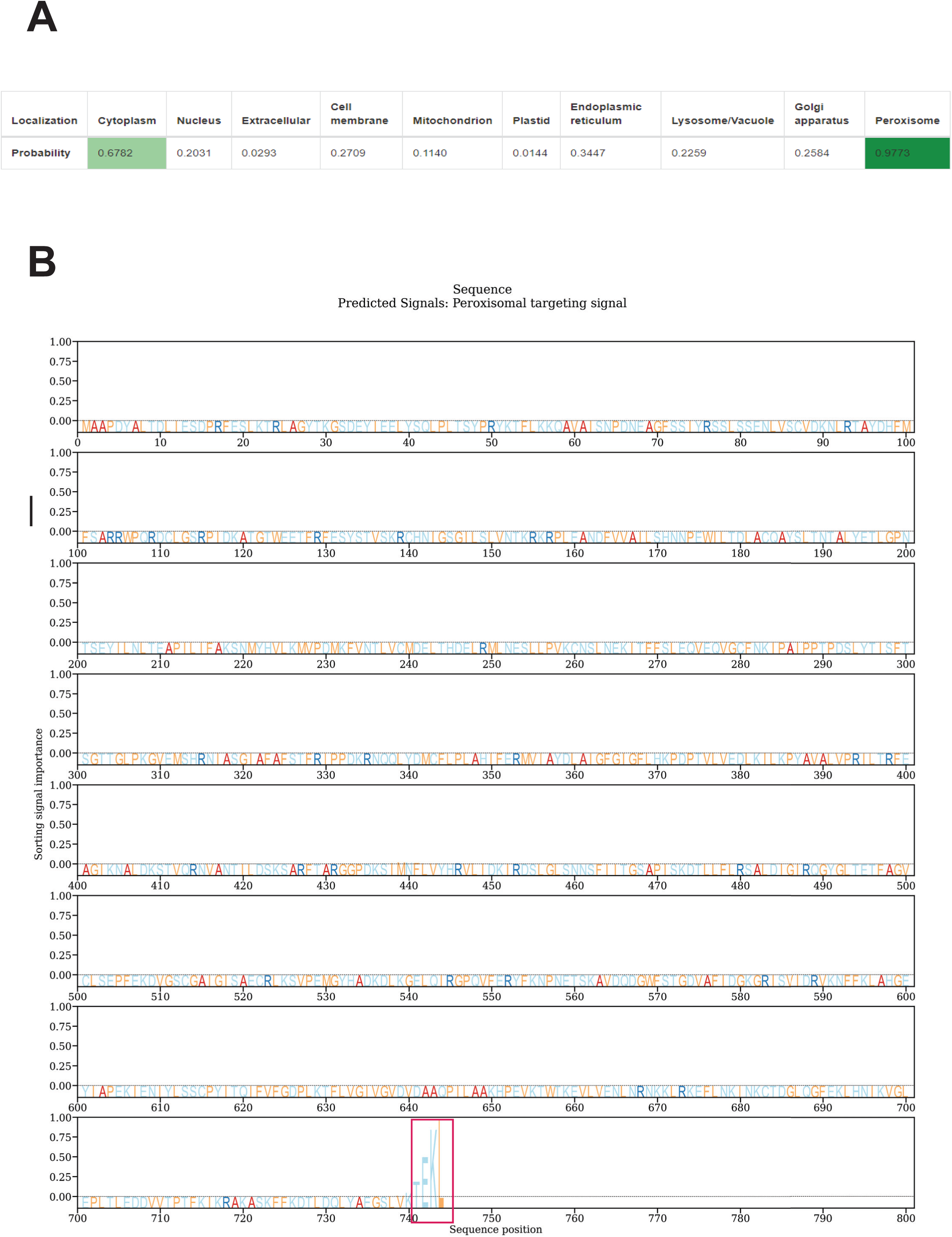
Prediction of the peroxisomal targeting sequence of Faa2 A) Probabilities of localization of Faa2 to different compartments was determined using the online tool DeepLoc2.0 Detection of the peroxisomal targeting sequence at the c-terminus of Faa2.

